# Strengthening Biofilms with Selective Metal Ions

**DOI:** 10.1101/2025.09.09.675226

**Authors:** Kiera J. Croland, R. Kōnane Bay

## Abstract

Biofilms are structured microbial communities consisting of bacteria embedded in a self-produced extracellular polymeric substance (EPS) that enables survival in diverse environments. The EPS integrates materials from the surrounding environment, such as metal ions, to provide additional mechanical protection to the embedded bacteria from environmental stressors. While previous studies demonstrated metal ions impact the erosion behavior of biofilms, key quantitative properties, such as yield stress, remain largely undocumented due to difficulties in handling these viscoelastic and soft biomaterials. In this work, we introduce a technique to characterize the impact of metal ions on the uniaxial stress-strain response of bacterial biofilms. Through applying this method to *Bacillus subtilis* pellicles, we demonstrate that the absorption of selective metal ions into the EPS significantly increases both the elastic modulus and yield stress, while decreasing failure strain. Notably, this effect can be reversed through the introduction of a strong chelating agent—while variations in pH alone have a negligible impact on measured mechanical properties. We compare our results to previous biofilm erosion studies and provide insights into how metal ions interactions can alter the mechanical behavior of biofilms, which will aid in future biofilm mitigation strategies for biofouling or healthcare applications.

## 1. Introduction

Biofilm-forming bacteria increase their resilience to the surrounding environment by aggregating together and encasing themselves in an extracellular polymeric substance (EPS). Biofilms are complex structures that readily colonize diverse surfaces, including human tissue,^[1,2]^ medical devices,^[3,4]^ and water distribution systems.^[5,6]^ The protective EPS provides constituent bacteria with structural integrity, surface adhesion, and protection from antimicrobial agents.^[7]^ As a result, biofilms pose significant challenges across many industries.^[5]^ In clinical settings, biofilm infections resist conventional antibiotic therapy and often require surgical intervention for complete removal.^[2]^ Biofilms that form on equipment surfaces in industrial settings, such as tanks and pipes, similarly require costly interventions through physical removal or equipment replacement.^[5,6]^ Despite these challenges, biofilms can be engineered to offer significant benefits. In wastewater treatment facilities, biofilms serve as bioreactors that efficiently degrade organic pollutants and remove contaminants from effluent streams.^[8]^ Additionally, biofilm properties can be exploited in engineered living materials (ELMs)^[9,10]^ to create stimuli-responsive intelligent systems. Across all applications, mechanical properties fundamentally govern biofilm behavior, yet characterization of these fragile biological materials remains technically challenging.^[11,12]^

Current approaches to biofilm mechanical characterization often yield results that are difficult to compare across systems due to a lack of standardization.^[11,12]^ Many mechanical testing methods for biofilms fail to preserve the structural integrity of biofilms (e.g., rheometry),^[1,13–19]^ do not yield quantitative mechanical parameters (e.g., no decipherable modulus,^[20]^ relative detachment values,^[13–15]^ stability^[21]^), and do not capture responses under physiologically relevant conditions.^[22–24]^ While microscale techniques (e.g., microrheology,^[25]^ AFM) preserve local structure, they do not reflect bulk mechanical behavior which is essential for understanding the macroscopic behavior of these inherently heterogenous structures. Furthermore, laboratory-grown biofilms often lack the complexity found in natural environments. *In situ* biofilms routinely incorporate external molecules or components from their surroundings into the EPS, altering material properties. Environmental factors, such as nutrients availability^[21]^ and host conditions,^[1]^ have been shown to influence EPS composition and production. Therefore, it is critical to understand how environmental stimuli, such as the exposure to metal ions^[13,14]^, biopolymers^[15]^, or antibiotics^[15]^, impact biofilm mechanics.

Metal ion interactions with the EPS is of particular interest due to their prevalence on colonized surfaces. Metal ions are abundant in biological settings, such as within the human body (e.g., blood, tissues) and on abiotic surfaces (e.g., metal pipes, industrial processing tanks). Additionally, bacteria are able to deteriorate metals through microbially induced corrosion (MIC).^[26]^ Although metal ions can exhibit toxicity to planktonic bacteria,^[13,27]^ biofilms can actively incorporate ions into the EPS leading to their long-term survival. Prior studies have demonstrated that treatment with metal ions can significantly influence biofilm mechanical properties, such as modulus (via shear rheology)^[13,14,19,28]^, adhesion or failure behavior (via erosion assays),^[13,14]^ and surface hydrophobicity.^[29]^ However, standard shear rheology techniques for biofilms often require scraping colonies from solid substrates and loading them onto a rheometer, which disrupts biofilm structure. Erosion studies preserve biofilm structure but do not yield quantitative mechanical parameters.

In this work, we introduce a new approach to tensile test bacterial biofilms. Unlike existing rheological and erosion approaches, tensile testing offers an alternative for probing biofilm mechanics, as it can capture both high-and low-strain responses and generate quantitative data. Additionally, tensile stresses are physiologically relevant to biofilms in flow environments, such as pipes or catheters, which experience tensile loading due to shear forces. We employ the soil-dwelling, non-pathogenic bacterium *Bacillus subtilis* (*B. subtilis*) as our model organism due to its well-characterized biofilm formation capabilities and precedence in biofilm mechanics research. *B. subtilis* is also ecologically and practically significant—it colonizes the human gastrointestinal tract,^[30]^ is used as a probiotic,^[31]^ and forms root-associated biofilms that promote nutrient uptake.^[32]^ Additionally, prior work has applied tensile testing *B. subtilis* colonies on agar plates^[20]^ as well as pellicles,^[22,23]^ which are biofilms which grow at the air-liquid interface; however, these studies were limited by long experimental timelines and ability to provide quantitative mechanical parameters, respectively. Our method utilizes a custom-built cantilever-based instrument, <u>T</u>he <u>U</u>niaxial <u>T</u>ensile <u>T</u>ester for <u>U</u>ltra<u>T</u>hin films (TUTTUT), which has previously been used for tensile testing on synthetic polymer ultrathin films (<200 nm).^[33–37]^ Critically, TUTTUT characterization preserves the native pellicle structure, enables the introduction of environmental cues at various points of biofilm development, and quantitatively measures mechanical properties across the complete strain range in one measurement. Using this method, we find that *B. subtilis* pellicles display viscoelastic behavior, consistent with previous reports.^[22,23]^ To investigate how environmental metal ion incorporation influences biofilm mechanical response, we characterized how *B. subtilis* pellicle stress-strain response changes upon exposure to Fe^3+^, Cu^2+^, Ca^2+^, and Na^+^. Our results suggest that metal ions can significantly impact elastic modulus (*E*), yield stress (σ_y)_, and failure strain (ε_f_), providing quantitative insight into metal ion effects on biofilm mechanics and demonstrating the versatility of our TUTTUT system for studying the impact of diverse environments on the mechanical behavior of biofilms.

## 2. Results and discussion

Here, we modify TUTTUT to directly measure the uniaxial stress-strain response of bacterial pellicles (**Figure 1A**). Prior to our measurements with biofilms, TUTTUT was used to characterize the stress-strain response of ultrathin polymer films (<200 nm).^**[33–37]**^ TUTTUT leverages a liquid support bath to manipulate difficult-to-handle materials, making it well-suited for characterizing ultrasoft biofilms (i.e.,∼100 Pa-1 kPa^**[13–15,22,23]**^). TUTTUT is a cantilever-based method, in which force resolution can be tuned by adjusting the material and geometry of the cantilever, enabling precise force measurements across a wide modulus range—from soft biofilms with elastic moduli in the Pa range to stiff polymer films in the GPa range. This makes TUTTUT broadly applicable to materials with widely varying mechanical properties. To the best of our knowledge, only one other research group has tested the tensile properties of pellicles.^**[22,23]**^ Their approach, however, relies on growing the biofilm on the liquid surface within their instrument, which limits the number of samples/variables that can be tested (∼100 biofilms in 5 years).

**Figure 1.**
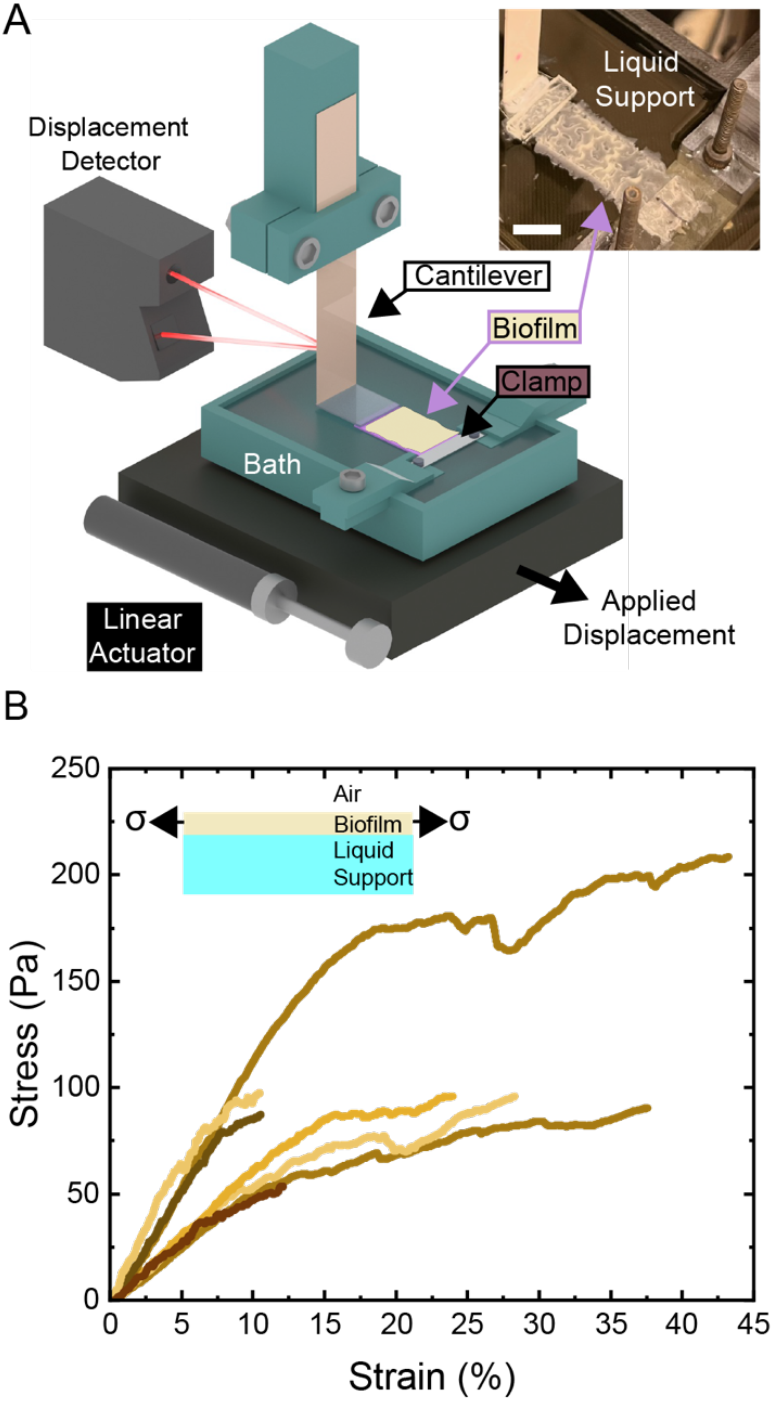
TUTTUT measurements of the complete uniaxial stress-strain response of *B. subtilis* pellicles. (A) Diagram of the uniaxial tensile tester for ultrathin films (TUTTUT) showing a pellicle atop the liquid support bath, held between a rigid boundary (clamp) and flexible cantilever. Inset (top right) shows a *B. subtilis* pellicle prior to tensile testing in TUTTUT (scale bar 1 cm). The pellicle is atop the liquid support bath, again, held between the cantilever and clamp. (B) Stress-strain curves for all untreated *B. subtilis* pellicles measured (n=7). Inset shows side view illustration of a pellicle being stretched on a liquid support.

In this work, *B. subtilis* pellicles are grown outside of the liquid support bath in TUTTUT (Figure 1A), allowing successive measurements without culture time restraints. We accomplish this by growing rectangular pellicles in 3D printed molds which generate two rectangular test samples from a connected media source (see Methods, Figure S1). After growth, the media or testing fluid (i.e., metal ion-supplemented or pH-adjusted media) is added to the culture container to float the biofilms out of the molds (Movie S1). One biofilm is transferred to TUTTUT, and the other is harvested onto a glass slide for determining the biofilm thickness (see Supporting Information). After transferring the biofilm to the bath, the biofilm is attached to the TUTTUT bath and the cantilever (Figure 1A). To perform tensile testing, a linear actuator stretches the biofilm at a constant strain rate (0.00295-0.00446 s^-1^) causing the cantilever to deflect. The cantilever is calibrated for force and displacement, and using the biofilm geometry, we calculate stress and strain (see Methods).

The stress-strain curves for untreated *B. subtilis* pellicles are shown in Figure 1B. Similarly to what has previously been observed,^**[22]**^ there is an initial linear-elastic regime followed by plastic deformation after a yield point. Interestingly, two distinct populations of stress-strain curves seem present for untreated pellicles; however, we found no clear factors (e.g. sample history or handling) that could account for the observed variation. We determine the elastic modulus, *E* ∼ 853 ± 314 Pa, the yield stress, σ_y_ ∼ 85.3 ± 44 Pa, and the failure strain, ε_f_ ∼ 25.9 ± 15.3 % *E* and σ _y_ are the same order of magnitude as the literature values (*E* ∼ 200-400 Pa and σ_y_ ∼ 50-200 Pa).^**[22,23]**^ Variations between our results and previously reported values may be attributed to differences in sample age and thickness measurement protocols. In this study, samples were tested on day 3, whereas prior studies evaluated samples aged for 2^**[22]**^ or 4-7^**[23]**^ days. Moreover, thickness was measured individually for each culture container in the present work, while previous studies assumed a constant average thickness of 350 μm across all samples.^**[22–24]**^ Furthermore, whereas previous researchers required five years to collect data on over 100 pellicles,^**[22]**^ TUTTUT enabled the characterization of 71 pellicles within just 22 testing days, corresponding to an average throughput of approximately ∼3.2 samples per day. This constitutes a substantial improvement in experimental efficiency.

Beyond increasing the throughput for measuring the mechanics of untreated pellicles, TUTTUT provides a platform to measure the influence of external environmental factors (e.g. free metal ions) on the mechanical properties of biofilms (Figure 2). Here, we assess the impact of Cu^2+^ and Fe^3+^, which are relevant to biofilms in copper and iron piping systems. Additionally, we probe the influence of Ca^2+^ and Na^+^, which are common free metal ions in water systems.^**[38,39]**^ After 3 days of growth, we expose pellicles to solutions of 50 mM metal ions of interest for one hour. Treatment with 50 mM Fe^3+^ and Cu^2+^ resulted in a pellicle color change from off-white to rust and blue (Figure 3), respectively, indicating the absorption of these ions into the biofilm matrix. For all metal ion treatments, we measure a similar stress-strain curve shape as the untreated biofilms (Figure 2A). Metal ion treatment leads to an increase in *E* and σ_y_ as well as a decrease in ε_f_ compared to the untreated biofilms (Figure 2B-D). To understand the impact of different metal ion treatments on biofilm mechanics, we compare *E*, σ _y_, and ε_f_.

**Figure 2.**
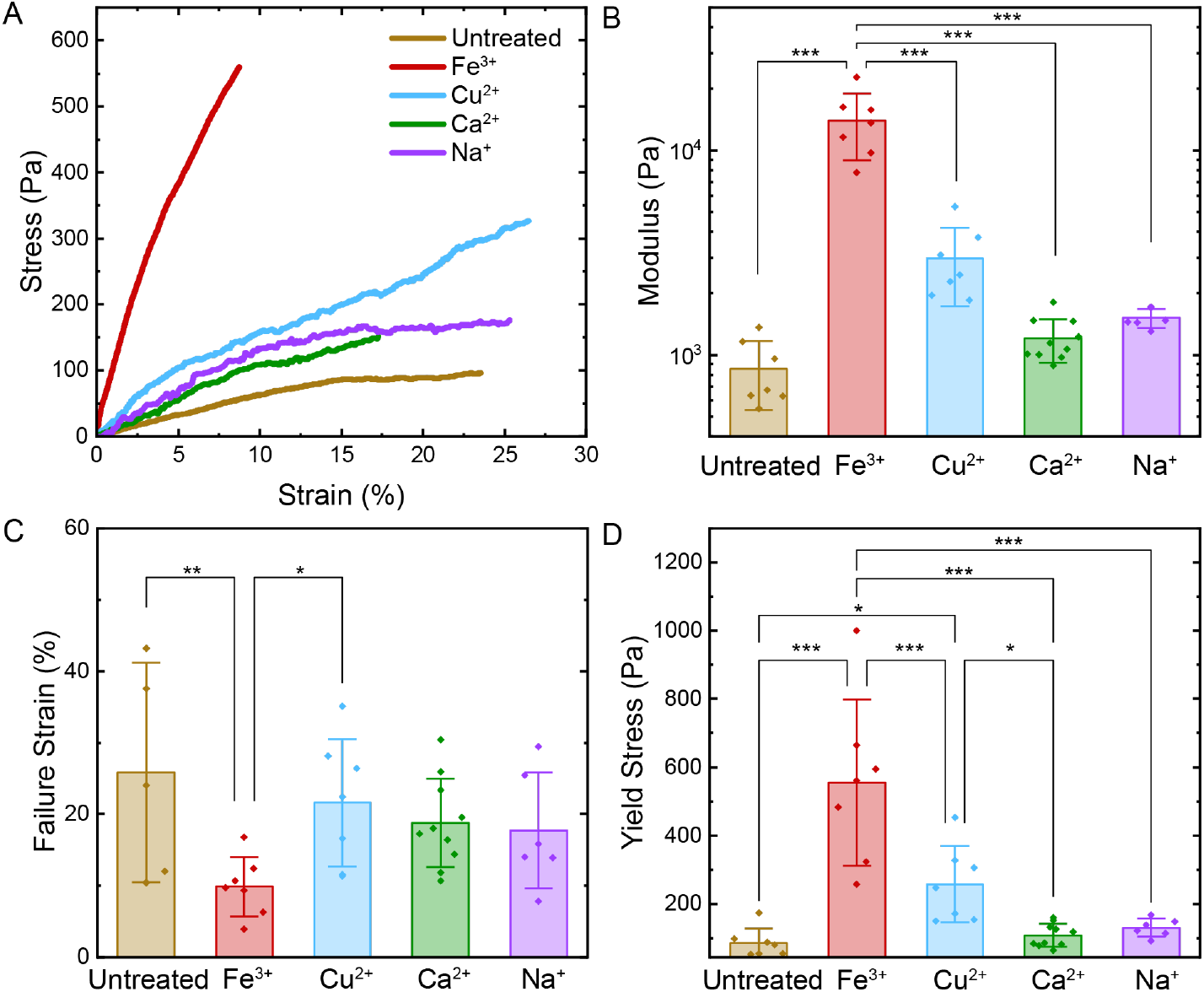
Treatment with selected metal ions (Fe^3+^, Cu^2+^, Ca^2+^, and Na^+^) impacts the stress-strain response of *B. subtilis* pellicles. (A) Individual representative stress-strain curves and extracted average (B) modulus, (C) failure strain, and (D) yield stress for untreated pellicles (n=7) as well as those incubated in media supplmented with 50 mM FeCl_3_ (n=7), CuSO_4_ (n=7), CaCl_2_ (n=10), or NaCl (n=6) for one hour. Data are represented as mean ± standard deviation (SD) (n≥3). Statistical significance is determined using a one-way Analysis of Variance (ANOVA) with a Fisher’s Least Significant Difference (LSD) post hoc test. p<0.05, p<0.01, and p<0.001 is reported with a *, **, and *** respectively.

**Figure 3.**
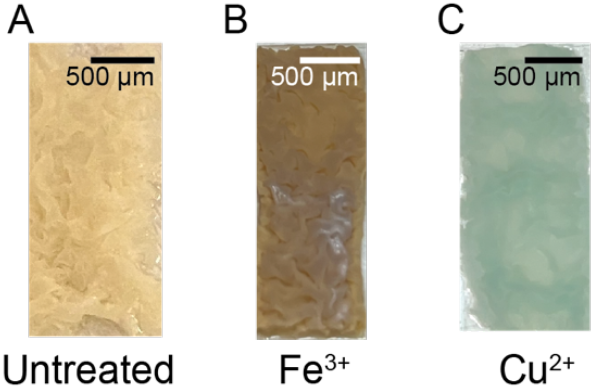
The color of (A) untreated biofilms displays significant changes following a one hour incubation with (B) FeCl_3_ or (C) CuSO_4_ indicating ion absorption into the biofilm EPS.

Pellicles treated with the trivalent ion, Fe^3+^ had the largest increase in the average *E*— an increase by an order of magnitude (Figure 2B). Pellicles treated with bivalent and monovalent ions exhibit a more moderate increases in *E*, although these changes are not statistically significant (Figure 2B). Researchers have observed similar increases in storage modulus (*G’*) for *Pseudomonas aeruginosa* (*P. aeruginosa*)^**[40]**^, *Escherichia coli*,^**[19]**^ and *B. subtilis* (B-1^**[13]**^ and NCIB 3610^**[14]**^) colonies with the addition of trivalent metal cations. Increases in *E* have also been observed in *P. aeruginosa* biofilms with Ca^2+^ exposure due to ionic crosslinking of alginate in the *P. aeruginosa* EPS.^**[41]**^ The increase in *E* with Fe^3+^ and Cu^2+^ treatment seen in this work is consistent with previously reported changes *G’* for ion-treated NCBI 3610 colonies;^**[14]**^ however, while we observe a slight, although not significant, increase in *E* with Na^+^ and Ca^2+^ treatment, previous measurements show a decrease or no change in *G’*.^**[13,14]**^ This discrepancy may arise from differences in sample preparation or growth method, which are known to influence measured mechanical properties for biofilms.^**[42]**^ Here, tensile tests are performed on pellicle biofilms, whereas previous work implemented shear rheology and erosion assays on biofilm colonies grown on agar plates.

The increase in *E* is associated with nonspecific ionic crosslinking between EPS components and metal ions. In *B. subtilis*, positively charged metal cations interact with negatively charged functional groups on proteins (e.g., carboxylic acid, aspartate), leading to increased crosslinking of the EPS.^**[26,43,44]**^ Additionally, cryo-electron microscopy on the *B. subtilis* EPS component TasA has suggested that aspartate residues on TasA can coordinate metal cations.^**[45]**^ Further, erosion assays performed on metal ion-treated NCIB 3610 colonies, specifically those performed on mutants lacking key EPS components, demonstrated that enhanced biofilm attachment arises from nonspecific interactions between the metal ions and the EPS components.^**[14]**^ Interestingly, this mechanism parallels that of metal-ion coordinated hydrogels, where reversible coordination between multivalent cations and polymer functional groups enhances mechanical properties, such as stiffness and toughness.^**[46]**^ Biofilms are often compared to colloidal hydrogels, where the cells are the colloids and the EPS is the crosslinked gel.^**[43,47,48]**^ Similar to the structural role that metal ion coordination plays in hydrogels, ion coordination in biofilms may reinforce the EPS through dynamic crosslinking interactions.

In further support of our ionic crosslinking hypothesis, we observe a trend of decreasing ε_f_ with the addition of the metal ions (Figure 2C). Although Fe^3+^ treated biofilms are the only group which displays a statistically significant decrease in ε_f_ from the untreated biofilms, a trend of decreasing ε_f_ with metal ion treatment is present. We propose that this decrease arises from an increase in network connectivity—specifically, higher crosslink density—which limits the ability of the biofilm to deform before failure. This interpretation aligns with findings from tensile tests on ionically-crosslinked hydrogels, where increasing crosslink density, achieved through increased metal cation content, leads to a reduction in ε_f_.^**[49]**^ In further support of ion-mediated stabilization, previous work showed supplementing *B. subtilis* NCIB 3610 cultures with Ca^2+^ limits biofilm dispersal by promoting structural stabilization,^**[50]**^ suggesting that metal ion-induced stabilization may influence biofilm persistence and failure. To our knowledge, this is the first quantification of ε_f_ in *B. subtilis* pellicles.

Two failure modes were commonly observed during tensile testing: either failure initiated at pre-existing cracks introduced during removal from culture molds or new cracks or holes formed and propagated under tension. While the presence of pre-existing cracks did not appear to significantly alter the ε_f_, samples with pre-existing cracks tended to exhibit lower maximum stress values at failure (Table S1). However, definitive conclusions cannot be drawn due to a limited number of replicates. We note that while ε_f_ does not correlate with previously reported erosion data for metal ion-treated biofilms,^**[14]**^ we do observe agreement when comparing to σ_y._

The addition of metal ions results in an increase in σ_y_, with Fe^3+^ producing the highest σ_y_, followed by Cu^2+^, Na^+^, and then Ca^2+^ (Figure 2D). We attribute these increases to ionic crosslinking, noting that the trend mirrors that observed increase in *E*—Fe^3+^ treatment produces the most pronounced increase, while Ca^2+^ results in the least; however, we note that the differences in σ_y_ between the untreated group and the Na^+^ and Ca^2+^ treated groups are not statistically significant. σ_y_ indicates resistance to irreversible deformation, making it relevant to understanding biofilm detachment behavior under shear flow or mechanical disruption. This may explain why biofilm detachment, as determined by erosion studies, appears to reflect the onset of yielding rather than complete mechanical failure.

Differences in σ_y_ values between metal ion treatments have several potential sources, including variations in the spatial distribution of the metal ions within the EPS or metal ion electronic structure. X-ray fluorescence (XRF) measurements support this idea, showing that Ca is distributed uniformly throughout the biofilm matrix, whereas Zn, Mn, and Fe are specifically enriched within biofilm wrinkles.^**[51]**^ The spatial distribution of ionic crosslinks can influence mechanical behavior by creating regions of localized stress concentration that are more susceptible to crack initiation,^**[52]**^ which can impact the measured σ_y_. In addition to spatial effects, differences in metal identity may also contribute to the observed changes in all properties measured. For example, in alginate hydrogels crosslinked with metal ions, *E* follows the trend, Fe^3+^ > Cu^2+^ > Ca^2+^, which is attributed to differences in ion charge, size, coordination number, and binding affinity. Trivalent ions, like Fe^3+^, can coordinate with three carboxylic groups on alginate, forming more crosslinks and generating stiffer networks. Divalent ions such as Cu^2+^ and Ca^2+^ typically coordinate with two groups, leading to more compliant structures.^**[53]**^ We observe a similar trend in biofilms, where Fe^3+^ treatment induces the largest change in measured properties among all ions tested. The difference in *E* for Cu^2+^ and Ca^2+^ crosslinkers in alginate hydrogels were attributed to the affinity of each ion to binding alginate.^**[53]**^ This could play a role in the differences in *E* observed in biofilms for divalent crosslinkers here: ions may have differing affinities for EPS polymers. Additionally, metal ions exhibit preferred coordination geometries based on their electronic configurations, but the specific binding environments presented by EPS components may constrain ions into less favorable geometries,^**[46]**^ giving rise to metal identity-dependent mechanical properties.

To directly test our ionic crosslinking hypothesis, we introduced an additional wash step to remove metal ions from pellicles pre-treated with Fe^3+^. Fe^3+^ was selected for this study because it caused the most pronounced mechanical differences from untreated biofilms. In this study, following treatment with 50 mM Fe^3+^, pellicles were subjected to a secondary wash step using LB-glycerol-manganese (LBGM) media or 25 mM ethylenediaminetetraacetic acid disodium salt (EDTA), a strong chelating agent. The LBGM media wash results in slight recovery of the stress-strain response (Figure 4A) and moderate reduction of the rust color associated with Fe^3+^ treatment (Figure 5). These changes suggest limited removal of Fe^3+^ ions and partial disruption of the ionic crosslinks within the EPS using LBGM media. In contrast, the EDTA wash produces a more dramatic color change, indicating more effective removal of Fe^3+^ from the EPS (Figure 5). Mechanically, the EDTA wash resulted in pellicle stress-strain curves and mechanical property metrics (*E*, σ_y_, or ε_f_) that closely resembled those of EDTA-only controls (i.e., pellicle treated with EDTA but no Fe^3+^), indicating reversal of Fe^3+^ induced stiffening (Figure 4B-D). The stronger impact of EDTA compared to the LBGM media wash is due to the high affinity of EDTA for metal ions,^**[54,55]**^ allowing it to more effectively disrupt Fe^3+^-EPS interactions. Trends seen here agree with erosion studies on *B. subtilis* B-1 colonies, where an EDTA wash following Fe^3+^ treatment restored biofilm detachment to ∼100%.^**[13]**^

**Figure 4.**
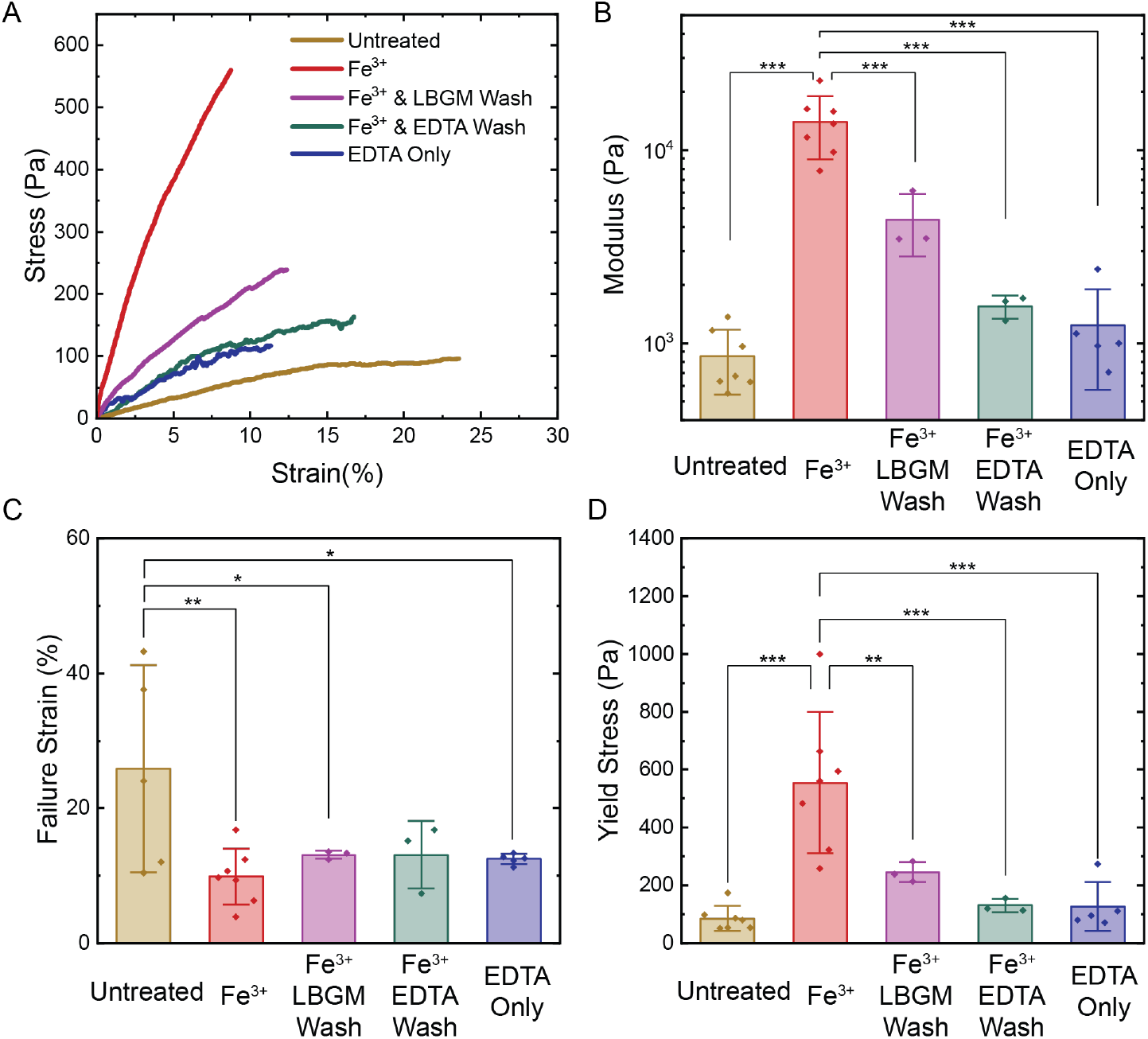
Mechanical characterization of pellicles following Fe^3+^ treatment and subsequent wash steps. Here, pellicles are exposed to 50 mM Fe^3+^ for one hour followed by a wash with 25 mM EDTA or LBGM media. (A) Representative stress-strain curves and comparison of average (B) elastic modulus, (C) failure strain, and (D) yield stress across treatment groups (untreated (n=7), Fe^3+^ (n=7), EDTA wash (n=3), LBGM wash (n=3), EDTA only (no Fe^3+^ pre-treatment, n=5)). Data are represented as mean ± SD (n≥3). Statistical significance is determined using a one-way ANOVA with a Fisher’s LSD post hoc test. p<0.05, p<0.01, and p<0.001 is reported with a *, **, and *** respectively.

**Figure 5.**
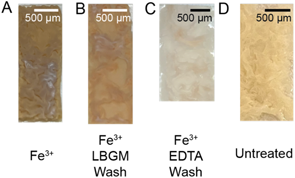
Representative images of a biofilm treated with (A) 50 mM Fe^3+^, pre-treated with 50 mM Fe^3+^ and exposed to a subsequent wash step with (B) LBGM media and (C) EDTA, and (D) an untreated biofilm.

Notably, treatment with EDTA alone seemed to impact biofilm mechanics. While the Na^+^ ions present in the EDTA solution could contribute to the observed changes, EDTA-specific interactions are also likely to play a role. In contrast to prior findings showing that EDTA compromises the structural integrity in biofilms formed by pathogenic bacteria,^**[58,59]**^ we see a slight decrease in ε_f_ for the EDTA only treatment, potentially indicative of increased network connectivity. This mechanical response could be influenced by interactions between EDTA and extracellular DNA (eDNA) in the EPS^**[56]**^ or pH effects.^**[51,57]**^ Previous studies have suggested that EDTA may enhance single-stranded DNA binding to *B. subtilis* cells,^**[56]**^ potentially introducing additional crosslinks within the biofilm matrix and contributing to the observed reduction in ε_f_. Additionally, previous work has shown that TasA fibers exhibit polymorphism and undergo configurational changes or aggregation in acidic conditions.^**[51,57]**^ This may result in biofilm matrix stiffening due to EDTA exposure and could impact stress-strain behavior.

Given the acidity of EDTA and that the addition of metal ions to LBGM media causes a reduction in pH (Table S2), we next explored the impact of pH on the biofilm tensile response. Similar to our metal ion studies, we perform TUTTUT testing on pellicles exposed to LBGM media adjusted to pH 2.0, 4.0, 8.0, and 10.0 for one hour (Figure 6, Figure S3). Treatment with pH adjusted media causes slight changes in measured mechanical properties for a few treatment groups, although much smaller than those caused by the metal ion solution with its corresponding pH. Thus, we attribute the increase in *E* and σ_y_, and decrease in ε_f_, to the presence of metal ions as opposed to the pH change. However, LBGM media adjusted to pH 4.0, which is closest to the pH of 25 mM EDTA, results in a moderate increase in *E* and σ_**y**_ and a decrease in ε_f_, suggesting that pH may contribute to the effects observed with EDTA treatment.

**Figure 6.**
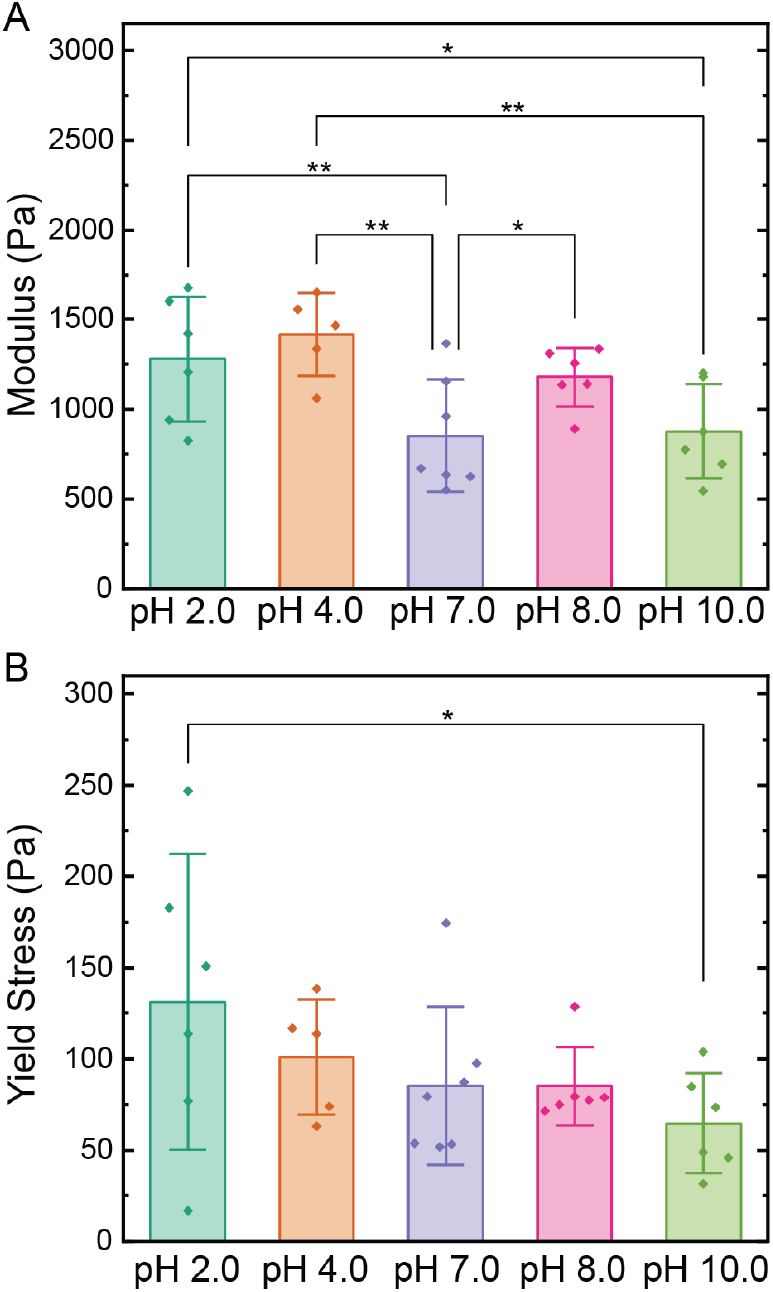
Treatment with LBGM media adjusted to pH 2.0 (n=5), 4.0 (n=5), 8.0 (n=6), and 10.0 (n=6) results in small changes in (A) average modulus, and (B) yield stress for *B. subtilis* pellicles. See Figure S3 for failure strain values and Table S2 for the pH of all treatment fluids used in this study. Data are represented as mean ± SD. Statistical significance is determined using a one-way ANOVA with a Fisher’s LSD post hoc test. p<0.05, p<0.01, and p<0.001 is reported with a *, **, and *** respectively.

## 3. Conclusion

We introduce a method, TUTTUT, to directly measure the uniaxial stress-strain relationship of bacterial biofilms. Using TUTTUT, we find that the *E*, ε_f_, and σ_y_ of *B. subtilis* pellicles are impacted by the absorption of the metal ions Fe^3+^, Cu^2+^, Na^+^, and Ca^2+^ into the biofilm matrix. pH is found to have a minimal influence on the observed changes in mechanical properties. Additionally, the mechanical properties of untreated pellicles can be partially recovered from Fe^3+^ pre-treated pellicles through the introduction of a chelating agent, EDTA. The role of metal ions in controlling biofilm mechanics highlights the importance of considering the ionic environment when developing strategies for biofilm removal or control. The presence of strongly bound metal ions, such as Fe^3+^, may confer increased mechanical stability, making biofilms more resistant to mechanical removal. Effective biofilm mitigation strategies may therefore require the identification and targeted removal of ions from the EPS. Furthermore, these results are relevant to a wide range of applications in which biofilms interact with materials—such as medical devices, water infrastructure, and industrial surfaces—where local ion availability may significantly influence biofilm structure, persistence, and response to treatment.

## 4. Experimental Section/Methods

### Bacterial culture

Liquid cultures of *B. subtilis* NCIB 3610 were generated by inoculating 2% w/v LB broth (Lennox) with a piece of frozen bacterial glycerol stock and incubating overnight at 30 °C and shaking at 240 rpm. Overnight cultures were then diluted 1000× in LB and are incubated at 30 °C and shaking at 240 rpm until an optical density (OD) at 600 nm of ∼0.1 was reached, resulting in our starting culture. To achieve the desired pellicle testing geometry, we 3D printed molds that have two rectangular holes (25 mm × 7.5 mm × 10 mm deep) and one circular hole for adding additional media to the mold after biofilm growth (Figure S1). Culture molds grow two pellicle strips from a connected media source: one is used for tensile testing and the other for thickness measurements. The molds were secured within sterile Magenta Plant Culture Boxes then filled with LBGM media (2% LB, 0.1 mM MnCl_2_, and 3% glycerol) such that there was ∼1 cm of media covering the bottom of the culture (7.5-35 mL). Our starting culture was inoculated into the molds, resulting in an OD_600_≈0.001. The culture box was covered with parafilm and incubated at room temperature for 3 days prior to mechanical testing.

### Biofilm harvesting and testing

After 3 days of growth, a scalpel was used to detach biofilm samples from the edges of the 3D printed molds. Biofilms were floated up out of their culture molds by adding additional media to the container (Video S1). Then, one biofilm was transferred from the culture container to TUTTUT bath. For metal ion studies, biofilm samples were floated in 50 mM FeCl_3_, CuSO_4_, CaCl_2_, or NaCl supplemented media for an exposure time of one hour prior to transferring to TUTTUT. The remaining biofilm was harvested on a glass slide for thickness measurements. Only biofilms in which both pellicle strips displayed a wrinkled morphology were used for measurements. Once in the TUTTUT bath, a clamp fabricated from rectangular coverglass slides (30 mm × 5 mm) with a ∼0.2 mm PDMS (1:10 cross-linker:silicone elastomer base) coating on one side was dropped onto one end of the biofilm and was secured into place on the bath using magnets. The free edge of the biofilm was then aligned with the extension piece of the cantilever, and the bath was raised to attach the sample to the cantilever grip. To perform tensile testing, a linear actuator on TUTTUT caused the clamp to move away from the cantilever at a fixed velocity (*v*_*actuator*_ = 0.1 mm s^-1^), stretching the biofilm at a fixed strain rate (0.00295-0.00446 s^-1^) and causing the cantilever to deflect (Movie S1). Tensile tests in which the pellicle slipped from either the clamp or cantilever grip were excluded from analysis.

### Thickness measurements

A Keyence VR-3000 3D Optical Profilometer (OP) was used to scan the 3D profiles of biofilms on glass slides. Using instrument analysis software, eight equally sized squares were selected within the sample, returning each square’s average thickness with respect to the instrument stage. The thickness of the glass slide was subtracted from measurements, and the eight thickness values were averaged to obtain average biofilm thickness for that culture container.

### Investigation of pH effects

LBGM media (pH 7.0) was prepared as described in Bacterial Culture and pH adjusted to 2.0, 4.0, 8.0, or 10.0 using 1M HCl and 1M NaOH. To determine the impact of pH on the biofilm stress-strain response, biofilms were cultured for 3 days, then floated in pH adjusted media for an exposure time of one hour prior to tensile testing.

### Chelation based analysis of biofilm-Fe^3+^ affinity

Pellicles were treated with 50 mM FeCl3 for one hour prior to transfer to a wash bath, containing either 25 mM EDTA or LBGM media. After one hour, biofilms were transferred to TUTTUT for mechanical testing as described in Biofilm harvesting and testing.

### Cantilever fabrication and calibration

Force readings were determined using the stiffness of cantilevers (*S*_*c*_) fabricated from rectangular polyethylene terephthalate (PET) sheets. Cantilever material was selected to be at least 2× the stiffness of biofilm samples. Cantilever calibrations and stress-strain calculations were performed as previously described.^[33]^ In brief, *S*_*c*_ was calculated by measuring resonance frequency (*f*) at varied cantilever lengths (*L*_*c*_). *f* was measured by a LK-G5000 Series Laser Displacement Sensor (1000 Hz) to capture the cantilever displacement (*δ*_*cant*_) that occurred in response to tapping the cantilever. A Fourier transform was fit to this data to determine *f*. Three *f* measurements were taken for five different *L*_*c*_ approximately 5 mm apart. We calculate the stiffness using *f, L*_*c*_, total cantilever length (*L*_*total*_), and total cantilever mass (*m*_*total*_) by 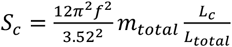 (Equation 1).^[60]^ To establish a relationship between laser displacement (*δ*_*laser*_) and *δ*_*cant*_, a displacement is applied in the x-direction using the linear actuator and the resulting δ_laser_ is measured. A linear fit yields slope *m*_*1*_. *δ*_*cant*_ is determined by dividing *δ*_*laser*_ by *m*_*1*_, and force is then determined by *F* = *S*_*c*_ ∗ δ_*cant*.)_. Then, biofilm displacement (*δ*_*film*_) is determined by δ_*film*_ = *v*_*actuator*_ ∗ *t* − δ_*cant*.)_, where *t* is time. Strain (*ε*) is determined by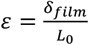, where *L*_*0*_ is the initial length of the film between grips. Stress (*σ*) is calculated by 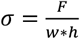, where *w* and *h* are biofilm width and thickness, respectively.

### Statistical analysis

All quantitative data are presented as mean ± standard deviation (SD) (n≥3). Statistical significance between sample means was determined using a one-way analysis of variance (ANOVA), followed by a Fisher’s Least Significant Difference (LSD) post hoc test for pairwise comparisons in OriginLab. Statistical significance with p<0.05, p<0.01, and p<0.001 is reported with a *, **, and *** respectively.

## Supporting information

Biofilm TUTTUT Method Video

## Acknowledgements

KC sincerely thanks Ava Crowley for supporting this work through generating diagrams of TUTTUT and for guidance and training in the Huli Materials Lab. We would like to acknowledge the Living Materials Laboratory of Wil Srubar at CU Boulder for providing the *B. subtilis* NCIB 3610 strain used in this work. We also thank the Joselle McCracken and the White laboratory at CU Boulder for supporting this work through providing use of the structured light profilometer for thickness measurements. This project was financially supported by the National Science Foundation Graduate Research Fellowship (NSF DGE # 2040434) and University of Colorado Boulder Startup Funds.

## Data Availability Statement

All elastic moduli, yield stresses, and failure strains extracted from stress-strain curves are available in the Supplemental Information. Stress-strain curves are available upon request.

