## Supplement information for "Strengthening Biofilms with Selective Metal Ions"

K. J. Croland & R. K. Bay

1. Department of Chemical and Biological Engineering, University of Colorado Boulder, Boulder, USA

2. Materials Science and Engineering Program, University of Colorado Boulder, Boulder, USA

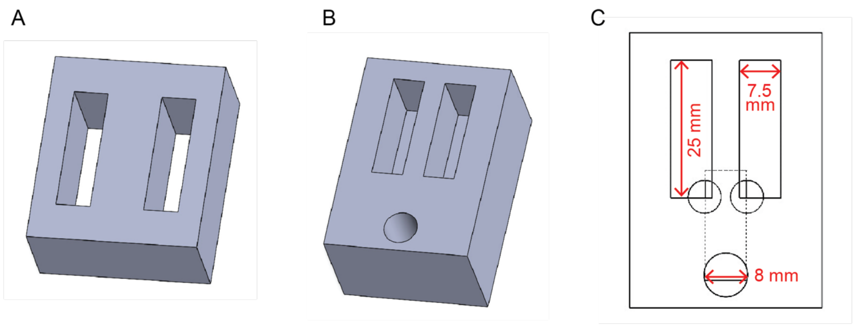


**Figure S1**. 3D renderings of pellicle culture molds used to generate rectangular biofilm strips for mechanical testing. Both molds have two 7.5 mm × 25 mm rectangular slits that define the sample culture regions. (A) Mold design which creates a pellicle within the mold region as well as throughout the cube culture container. Containers are filled with 35 mL of media, and pellicles form both inside the slits and across the liquid surface covering the culture container; only the in-mold pellicles are used, while those forming externally are sacrificed. Pellicles are floated from the mold by adding fluid to the sacrificed region to apply upward pressure beneath the culture slits. (B) Alternative mold design with an integrated media reservoir and a central media addition hole (shown in Movie S1). This version is filled with 7.5 mL of media directly into the mold. For testing, the pellicle that forms over the addition hole is first removed, and fresh media is added through the opening to float the biofilm strips from below using the same fluid pressure approach as in (A). (C) Internal view of the mold shown in (B), showing the fluid channel geometry and media addition port.



**Commented [RB1]:** Can you put the dimensions on the figure C?

**Commented [KC2R1]:** Done

**Table S1.** Average modulus (Pa), failure strain (%), maximum stress at failure (Pa), and yield stress (Pa) for untreated B. subtilis pellicles as well as those treated with 50 mM $FeCl_3$, $CuSO_4$, $CaCl_2$, or NaCl for 1 hour. Data is filtered and presented as an average (if n ≥2) standard deviation (if n≥3) for the presence of a pre-existing crack (if n=1, recorded values represent a single data point). Cracks often initiated during pellicle removal and transfer from culture molds.

**Commented [KB3]:** Can you put the standard error too? for samples that have more than 2?

**Commented [KC4R3]:** addressed

| Treatment | Pre-existing crack? (Y/N) | # samples (n) | Modulus (Pa) | Failure strain (%) | Failure stress (Pa) | Yield Stress (Pa) |
|---|---|---|---|---|---|---|
| Untreated | N | 6 | 768±239 | 28.4±15.1 | 124±67.1 | 83.3±47.1 |
| | Y | 1 | 1370 | 10.4 | 97.5 | 97.5 |
| $Fe^{3+}$ | N | 6 | 15000±4590 | 9.76±4.53 | 666±225 | 605±226 |
| | Y | 1 | 7800 | 10.7 | 305 | 258 |
| $Cu^{2+}$ | N | 5 | 3380±1220 | 18.8±8.02 | 339±213 | 271±123 |
| | Y | 2 | 1910 | 28.7 | 312 | 228 |
| $Ca^{2+}$ | N | 5 | 1150±223 | 21.3±6.70 | 130. ±33.7 | 116±34.2 |
| | Y | 4 | 1270±362 | 16.2±5.16 | 108±37.0 | 100. ±34.9 |
| $Na^{+}$ | N | 6 | 1520±166 | 17.8±8.08 | 142±29.2 | 130. ±27.0 |
| | Y | 0 | N/A | N/A | N/A | N/A |

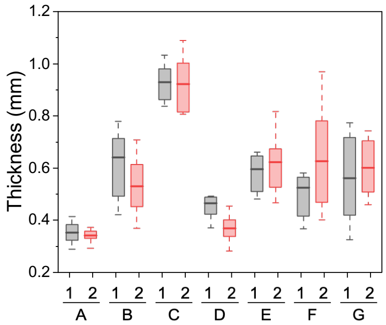


**Figure S2**. Comparison of the thickness of the two pellicle strips generated from the same culture mold. Pellicles are grown within culture molds shown in Figure S1, and instead of sacrificing one pellicle strip for tensile testing (as is done for all tensile tests in this work), the thickness of both strips is measured to validate the assumption that the two pellicle strips from a single culture mold are the same thickness. Letters (A-G) represent different culture molds and numbers (1 or 2) represent the 2 samples from the same culture mold. Boxes represent the range from 25%-75%, whiskers represent the range within 1.5 times the interquartile range (IQR), and bolded lines represent the mean. We find that pellicles from the same culture mold are 10.6 ± 8.88 % different from one another, on average.

**Table S2**. pH of media and all treaments

| Treatment | pH |
|---|---|
| LBGM Media | 7.0 |
| 50 mM $FeCl_3$ in LBGM | 2.1 |
| 50 mM $CuSO_4$ in LBGM Media | 3.5 |
| 50 mM $CaCl_2$ in LBGM Media | 6.5 |
| 50 mM NaCl in LBGM Media | 6.9 |
| 25 mM EDTA | 5.1 |

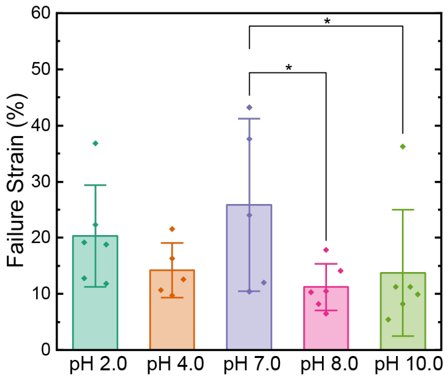


**Figure S3**. Failure strain for *B. subtilis* pellicles treated with LB-glycerol-manganese (LBGM) media adjusted to pH 2.0 (n=5), 4.0 (n=5), 8.0 (n=6), and 10.0 (n=6). Data are represented as mean ± standard deviation ($n \geq 3$). Statistical significance was determined using a one-way ANOVA with a Fisher's Least Significant Difference (LSD) post hoc test. $p<0.05$, $p<0.01$, and $p<0.001$ is reported with a *, **, and *** respectively.
